# Modulation of all-*trans* retinoic acid by light and dopamine in the murine eye

**DOI:** 10.1101/2024.12.06.627245

**Authors:** Sarah Talwar, Reece Mazade, Melissa Bentley-Ford, Jianshi Yu, Nageswara Pilli, Maureen A Kane, C Ross Ethier, Machelle T Pardue

**Author notes:** Corresponding author: Machelle T Pardue 1365B Clifton Road NE Atlanta, GA 30322. co-first authors (Sarah Talwar, Reece Mazade). **Funding Statement** NIH R01 EY016435 (MTP), NIH R01 EY033361 (MTP, CRE, MAK), NIH F32 EY035573 (MBF), NIH T32 EY007092 (MBF), Dept. of Veterans Affairs Research Career Scientist Award RX003134 (MTP), Research to Prevent Blindness Challenge Award (Emory Ophthalmology), Georgia Research Alliance (CRE). **Disclosures** The authors do not have any commercial relationship disclosures related to this work.

## Abstract

**Purpose:** Ambient light exposure is linked to myopia development in children and affects myopia susceptibility in animal models. Currently, it is unclear which signals mediate the effects of light on myopia. All-*trans* retinoic acid (atRA) and dopamine (DA) oppositely influence experimental myopia and may be involved in the retino-scleral signaling cascade underlying myopic eye growth. However, how ocular atRA responds to different lighting and whether atRA and DA interact remains unknown.

**Methods:** Dark-adapted C57BL/6J mice (29-31 days old) were exposed to Dim (1 lux), Mid (59 lux), or Bright (12,000 lux) ambient lighting for 5-60 minutes. Some mice were also systemically administered the DA precursor, LDOPA, or atRA prior to light exposure. After exposure, the retina and the back-of-the-eye (BOE) were collected and analyzed for levels of atRA, DA, and the DA metabolite, DOPAC.

**Results:** DA turnover (DOPAC/DA ratio) in the retina increased in magnitude after only five minutes of exposure to higher ambient luminance but was minimal in the BOE. In contrast, atRA levels in the retina and BOE significantly decreased with higher ambient luminance and longer duration exposure. Intriguingly, LDOPA-treated mice had a transient reduction in retinal atRA compared to saline-treated mice, whereas atRA treatment had no effect on ocular DA.

**Conclusions:** Ocular atRA was affected by the duration of exposure to different ambient lighting and retinal atRA levels decreased with increased DA. Overall, these data suggest specific interactions between ambient lighting, atRA, and DA that could have implications for the retino-scleral signaling cascade underlying myopic eye growth.

## Introduction

Outdoors, the eye is exposed to gradual changes in ambient brightness of up to 10 log units from night to day^1^. However, children spend most of their time indoors, which has a significantly restricted luminance range and reduced luminance intensity compared to daytime outdoor environments (>2 log units)^2–4^. Time spent indoors is thought to be a primary driver for the increased prevalence of myopia, or nearsightedness, worldwide^5^. Myopia results from altered eye growth during childhood and is characterized by an elongated eyeball where the visual image focuses in front of the retina, leading to blurred vision. While blurred vision can be optically corrected, myopia cannot be reversed, and myopic eyes have an increased risk of developing blinding complications^6, 7^. Therefore, understanding the mechanisms that drive myopia development is critical to developing effective interventions. Unsurprisingly, ambient light level, as one of the major differences between indoor and outdoor environments, has pronounced effects on myopia development.

Increasing ambient brightness during the critical period of ocular growth attenuates experimental myopia across animal models^8–17^. Likewise, children who spend more time in bright outdoor settings have reduced onset of myopia^18–22^, and emmetropic children spend more time in brighter environments than myopic children^3, 18^. Interestingly, recent work suggests that dim lighting (e.g. nighttime levels) is also protective in children and myopic mice, while mid-light conditions (e.g. those more often experienced indoors) are associated with higher levels of myopia^3, 17^. However, the specific signaling mechanisms that underlie the effect of light level on myopia development are still not fully understood.

Myopigenesis likely requires that visual cues, such as ambient light level, initiate or modulate a retino-scleral signaling cascade mediated by multiple signaling molecules. Two candidate molecules that are associated with both myopia and ambient light levels are dopamine (DA) and all-*trans* retinoic acid (atRA). DA is considered a “stop” signal for ocular growth and protects against myopia development^23, 24^ while atRA is associated with myopia progression (^25, 26^, for a full review see^27^). Furthermore, retinal DA is released from dopaminergic amacrine cells (DACs), increases with light, reaches peak steady state levels within 15 minutes, and is highest during the daytime^28–30^. Additionally, DA turnover (ratio of the DA metabolite 3,4-dihydroxyphenylacetic acid (DOPAC) to DA) increases with ambient light level in mice after chronic (2 weeks), prolonged (1- 3 hours), or shorter (15 minutes, isolated eye cups) exposure^17, 30^. Likewise, atRA, generated in multiple ocular tissues^31–39^, increases in isolated dark-adapted mouse retina in response to light^34^.

Despite these findings, two critical questions remain. First, ambient brightness is dynamic and can rapidly change (e.g. order of minutes) due to changes in artificial lighting indoors or eye movements in the visual scene^40^. However, it is unknown how atRA (or DA) levels change in the eye within this timeframe after exposure to light levels common in daily life (e.g. 1 – 10,000 lux). Second, it is not well-understood if DA interacts with atRA since they have opposite associations with myopia and luminance. Here, we answer these questions by briefly exposing mice to different ambient lighting before measuring levels of DA and atRA in the **retina** and “back of the eye” (**BOE**: whole eye minus the lens and retina). Furthermore, we investigate potential interactions between DA and atRA during acute light exposure.

## Methods

### Animals and experimental paradigms

C57BL/6J mice (Jackson Labs, Bar Harbor, ME, USA) were housed in a 12:12hr light:dark cycle in standard animal facility lighting (30-240 lux in a standard cage) at the Atlanta Veterans Affairs (VA) Healthcare System or Emory University School of Medicine. All procedures were approved by the Atlanta VA or Emory University Institutional Animal Care and Use Committee and adhered to the ARVO Statement for the Use of Animals in Ophthalmic and Vision Research.

In experiment 1, postnatal age (P) 29-31 mice were dark adapted for two hours before being exposed to Dim (1 lux, 0.07 µW/cm^2^, 11.27 log photon flux), Mid (59 lux, 3.57 µW/cm^2^, 12.99 log photon flux), or Bright (12,000 lux, 895.03 µW/cm^2^, 15.40 log photon flux) ambient lighting for 5, 15, or 60 minutes (**Figure 1A**). The Dim, Mid, and Bright levels were chosen to mimic the levels associated with dimmer indoor environments and brighter outdoor environments (**Figure 1B-C)**. During the experiment, mice were only exposed to ambient light during their exposure time; otherwise, their cages were protected from light. To expose animals to light, individual mice were placed in a clean, empty cage, without the cage top or food hopper, after which the cage was placed directly under the light source. This provided uniform light exposure and prevented shadows. Ambient lighting for Dim and Mid light levels was controlled using light-tight LED cabinets (Actimetrics, Wilmette, IL, USA) while daylight LED light panels (5,600K, Genaray SpectroLED Studio 800 Bi-Color, USA) were used for Bright light levels. Before each experiment, illuminance (lux) was measured at the floor of an empty cage using a lux meter (VWR Traceable Dual-Range, Radnor, PA, USA) to ensure the appropriate light level. Light spectrograms were measured (Sekonic Spectromaster C-7000, North White Plains, NY, USA) and transformed into irradiance and log photon flux based on mouse retinal sensitivities using the Rodent Irradiance Toolbox^41, 42^ (**Figure 1B-C)**. For consistency across experiments, all exposures and collections were performed between 13:00 and 15:00. Pupil size was not artificially controlled (e.g. via dilation) to model natural physiological visual states in different ambient lighting. Ocular levels of DA (male and female mice) and atRA (male mice) were measured in tissue collected immediately following light exposure. A total of 145 mice were used for experiment 1.

**Figure 1.**
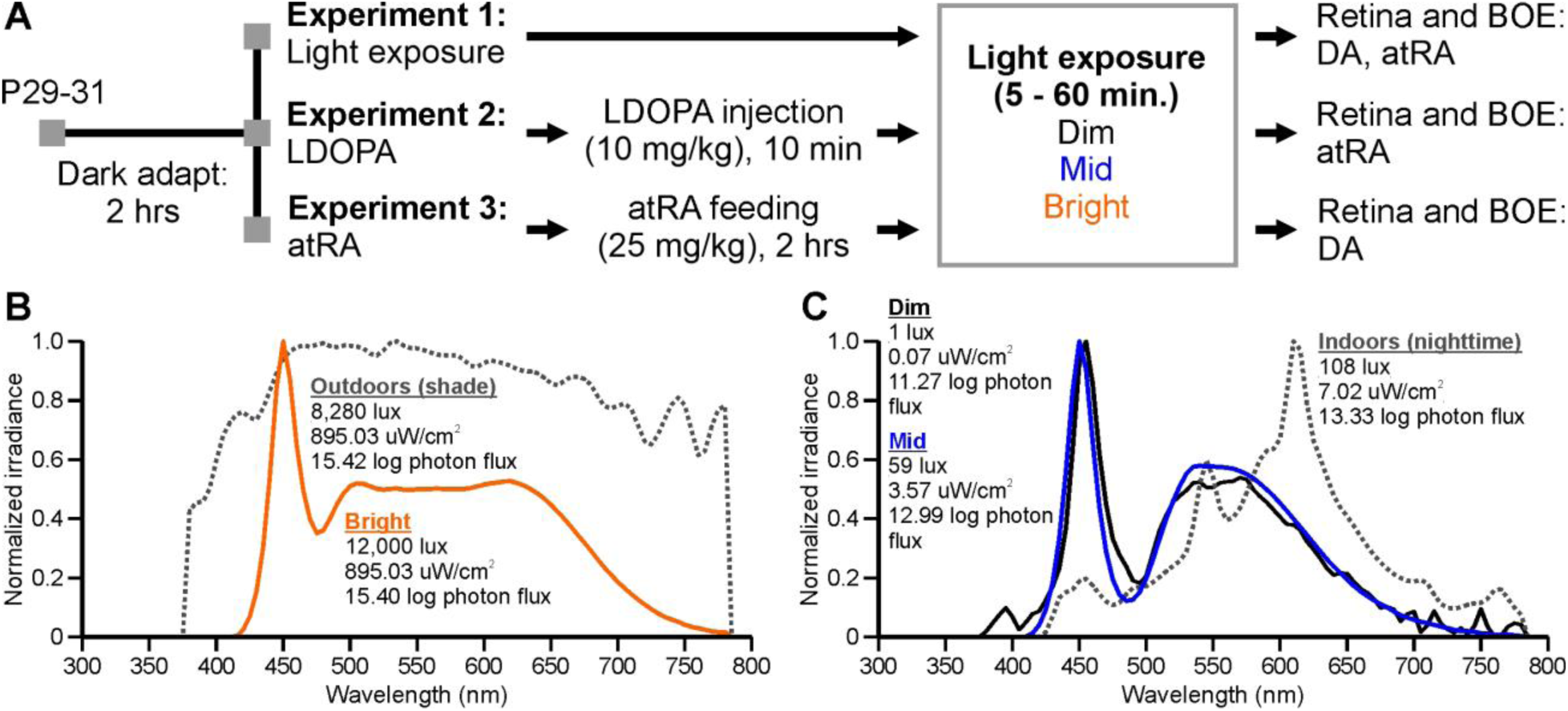
Experimental paradigms and light exposure. A. Mice, P29-31, were dark adapted for 2 hours before exposure to Dim, Mid, or Bright ambient lighting for 5, 15, or 60 minutes. In separate experiments, mice were given L-3,4-dihydroxyphenylalanine (LDOPA) or atRA at 10 minutes or 2 hours prior to light exposure, respectively. DA and atRA levels were measured from retinal or “back of the eye” (BOE: whole eye minus the lens and retina) collections. B-C. Normalized spectra and intensity values (human lux and mouse-adjusted log photon flux) of the Dim, Mid, and Bright experimental LED light sources compared to outdoor sunlight (B) or indoor artificial lighting (C).

In experiment 2 (**Figure 1A**), the influence of DA on ocular atRA levels was tested. Male and female mice received an intraperitoneal injection (IP) of L-3,4-dihydroxyphenylalanine (LDOPA) (dose: 10 mg/kg), a precursor of DA, 10 minutes before Mid light exposure as described for experiment 1. LDOPA (L-Dopa-(phenyl-d3, Sigma-Aldrich, Saint Louis, MO, USA) was dissolved in 0.9% sterile saline and administered in the dark under dim red light. Control mice were given 0.9% saline IP injections. After light exposure, retina and BOE samples were immediately collected to measure atRA levels. In a subset of mice, retina and BOE levels of DA were also measured after exposure to Bright lighting to serve as a positive control. A total of 48 mice were used for experiment 2.

In experiment 3 (**Figure 1A**), the influence of atRA on ocular DA was tested. Male mice were trained to voluntarily eat sugar icing pellets. First, mice underwent a training period for three days with plain sugar pellets to ensure they would consume the pellet during the experiment. atRA pellets consisted of sugar icing (sugar, water, corn syrup; Betty Crocker Cookie Icing, white) mixed with 1% atRA (≥98% pure, Cat # R2625; Sigma-Aldrich, Saint Louis, MO, USA) by weight. Vehicle pellets contained only sugar icing. The day before their use, atRA pellet formation was performed quickly in dim red light by mixing atRA directly into sugar icing because atRA isomerizes in light^43, 44^. Afterwards, individual pellets of the mixture were created, placed in light-tight boxes, and allowed to set overnight at 4°C. Vehicle pellets were formed and allowed to set overnight at 4°C.

After appropriate training and acclimation^26^, mice were weighed and placed in individual empty cages and given either a vehicle or an atRA pellet (dose: 25 mg/kg). Mice were dark adapted for two hours while they ate the pellet. The cages were frequently checked using a dim red light to note the time it took the mice to consume the pellets. Two hours after consuming the pellet, mice were exposed to Dim, Mid, and Bright ambient light levels as described for experiment 1. Following light exposure, retina and BOE samples were immediately collected for DA measurements. A total of 48 mice were used for experiment 3.

### Retina and BOE extraction

Retina and BOE samples were collected under the same light illumination as the exposure. After ambient light exposure, mice were euthanized via cervical dislocation, and the retina and BOE were extracted from the right (OD) and left (OS) eyes. First, the retina was extracted from the globe by squeezing the back of the eye through a slit in the cornea in one fluid motion to avoid contamination by the retinal pigment epithelium. The BOE was then extracted rapidly to minimize extra-ocular tissue and optic nerve contamination. The total time for sample collection (retina and BOE from both eyes) was <2 minutes. For consistency, the ocular samples from the OD eye were always extracted first. Both the retina and BOE samples were immediately snap frozen on dry ice and stored at −80°C. For DA measurements, OD and OS eye samples were collected separately. For atRA analysis, OD and OS eye samples were combined.

### Analysis of tissue samples

#### DA analysis

Samples were analyzed by high-performance liquid chromatography (HPLC) for both DA and its metabolite, DOPAC. Samples were homogenized in 0.1M perchloric acid, sonicated (30% duty cycle), and centrifuged at 4°C. The supernatant was then centrifuge-filtered and delivered to the Emory HPLC Bioanalytical Core facility for monoamine analysis using an ESA 5600A CoulArray detection system, equipped with an ESA Model 584 pump and an ESA 542 refrigerated autosampler, as described previously^45^. Briefly, separations were performed using a Hypersil 150 × 3 mm (3 uM particle size) C18 column at 28°C. The mobile phase consisted of 8% acetonitrile, 75 mM NaH2PO4, 1.7 mM 1-octanesulfonic acid sodium, and 0.025% trimethylamine at pH 3.0. The samples were eluted isocratically at 0.4 mL/min and detected using a 6210 electrochemical cell (ESA, Bedford, MA) equipped with 5020 guard cell. Guard cell potential was set at 500 mV, while analytical cell potentials were −175, 150 and 350 mV. The analytes were identified by the matching criteria of retention time to known standards of DA and DOPAC (Sigma-Aldrich, Saint Louis, MO, USA). Compounds were quantified by comparing peak areas to those of standards on the dominant sensor. Individuals performing DA analysis were blinded to the treatment groups. Total protein of each sample was determined using the Pierce bicinchoninic acid (BCA) assay. Samples were normalized by dividing the DA and DOPAC HPLC concentrations by the total protein (average of triplicates) to reduce variability due to differences in sample sizes. DA and DOPAC levels are presented as ng per mg of tissue. Dopamine turnover was calculated as the DOPAC/DA ratio before normalizing to total protein.

#### Retinoid analysis

Retina and BOE samples from OD and OS eyes were combined due to small tissue quantity for retinoid detection (mean ± standard deviation sample mass; retina: 5.14 ± 1.55 mg; BOE: 8.13 ± 1.89 mg). Snap-frozen samples were stored at −80 °C before they were shipped from Atlanta, Georgia to Baltimore, Maryland (Kane Laboratory) on dry ice for liquid chromatography - tandem mass spectrometry (LC-MS/MS) retinoid analysis. atRA was extracted following a two-step liquid-liquid extraction under dim yellow light, as previously described^46^. Briefly, after homogenizing, extracting, and facilitating phase separation of the tissue samples, the organic phase containing retinol and retinyl esters was isolated. Retinol and retinyl esters were quantified by Waters Acquity UPLC H-class system with a UV detector^47^. The remaining aqueous phase of the tissue homogenates was acidified and extracted, and the hexane layer was removed, containing atRA. This hexane layer was further processed before LC-MS/MS was performed with a Shimadzu Prominence UFCL XR liquid chromatography system (Shimadzu, Columbia, MD) coupled to an AB Sciex 6500 QTRAP hybrid triple quadrupole mass spectrometer (AB Sciex, Framingham, MA) using atmospheric pressure chemical ionization (APCI) operated in positive ion mode as described previously^43, 44, 48, 49^. The retinoid content in each sample was normalized to the tissue weight. atRA levels are presented as pmol per gram of tissue. Retinol and retinyl esters are presented as nmol per gram of tissue. Individuals performing atRA analysis were blinded to the treatment groups.

### Statistics

Statistical analysis was conducted using GraphPad Prism 8 (San Diego, CA, USA). For DA and DOPAC analysis, OD and OS eye measurements were averaged together. If a sample from one eye could not be collected, then only one sample was included. For the BOE, DOPAC levels in many samples were below the HPLC detection threshold of 0.1 ng/ml. Therefore, to include these valuable samples in the data set, the DOPAC concentration was arbitrarily set to 0.1 ng/ml before normalization as noted in the figures.

In experiment 1, the effect of light levels was tested using a one-way ANOVA with Tukey’s post hoc multiple comparisons test. In experiments 2 and 3, the effects of light level and LDOPA, light level and exposure duration, or light level and atRA were tested using two-way repeated measures ANOVAs. If interaction effects were found to be significant, Tukey’s post hoc multiple comparisons test were conducted. For all datasets, ROUT outlier tests were performed (Q=5%, GraphPad Prism 8), and any outliers were excluded. Note that retinol and retinyl ester measurements were subject to possible batch effects, potentially due to changes in dietary levels of vitamin A in mouse chow (Jackson Labs), and therefore the data is shown in tables with standard deviations. Sample sizes are noted in the figure captions or in the Results section. Data are presented as mean ± SEM, unless otherwise noted, and significance is denoted as *p<0.05, **p<0.01, and ***p<0.001.

## Results

### The increase in retinal DA turnover with light intensity is apparent after acute light exposure

Retinal DA and DA turnover (DOPAC/DA ratio) are strongly linked to light levels. Previous reports demonstrate that DA turnover in mice increases with ambient light intensity after more prolonged light exposure (tens of minutes to days)^17, 30^. Despite DA reaching steady-state levels by 15 minutes after light onset^30^, it is unclear whether DA turnover is affected after even briefer exposure to a wide range of ambient lighting. Therefore, mice were exposed to Dim, Mid, and Bright ambient light (see Methods) for 5 minutes, after which retina and BOE samples were collected to measure levels of DA and DOPAC and to calculate DA turnover. Retinal DA levels were highest after exposure to Dim and Mid light, yet significantly decreased after Bright light exposure (**Figure 2A**; Dim: 4.70±0.28 ng/mg, Mid: 4.67±0.18 ng/mg, Bright: 3.94±0.17 ng/mg, one-way ANOVA, effect of light: p=0.015; Dim vs Bright: p<0.05, Mid vs Bright: p<0.05). As a positive control, a subset of mice was treated with IP injections of LDOPA (10 mg/kg) and exposed to Bright ambient lighting. Retinal DA levels were significantly elevated after LDOPA delivery compared to all light levels (8.50±0.76 ng/mg, p<0.001). In contrast, retinal DOPAC levels increased with light intensity (**Figure 2B**). DOPAC levels were lowest after Dim light exposure (0.56±0.06 ng/mg) but significantly increased after Mid (0.99±0.08 ng/mg) or Bright light exposure (0.99±0.08 ng/mg, one-way ANOVA, effect of light: p<0.001; Dim vs Mid: p<0.01, Dim vs Bright: p<0.001). As with DA, LDOPA-treated mice had significantly elevated retinal DOPAC levels (4.33±0.37 ng/mg, p<0.001). Due to the opposing effects of acute ambient light exposure on retinal DA and DOPAC, DA turnover, calculated as the DOPAC/DA ratio, significantly increased after exposure to bright ambient light (**Figure 2C**, Dim vs Mid vs Bright, 0.12±0.01 vs 0.19±0.01 vs 0.25±0.02, one-way ANOVA, effect of light: p<0.001; Dim vs Mid: p<0.01, Dim vs Bright: p<0.001, Mid vs Bright: p<0.05). Likewise, LDOPA treatment doubled DA turnover (0.52±0.03, p<0.001).

**Figure 2.**
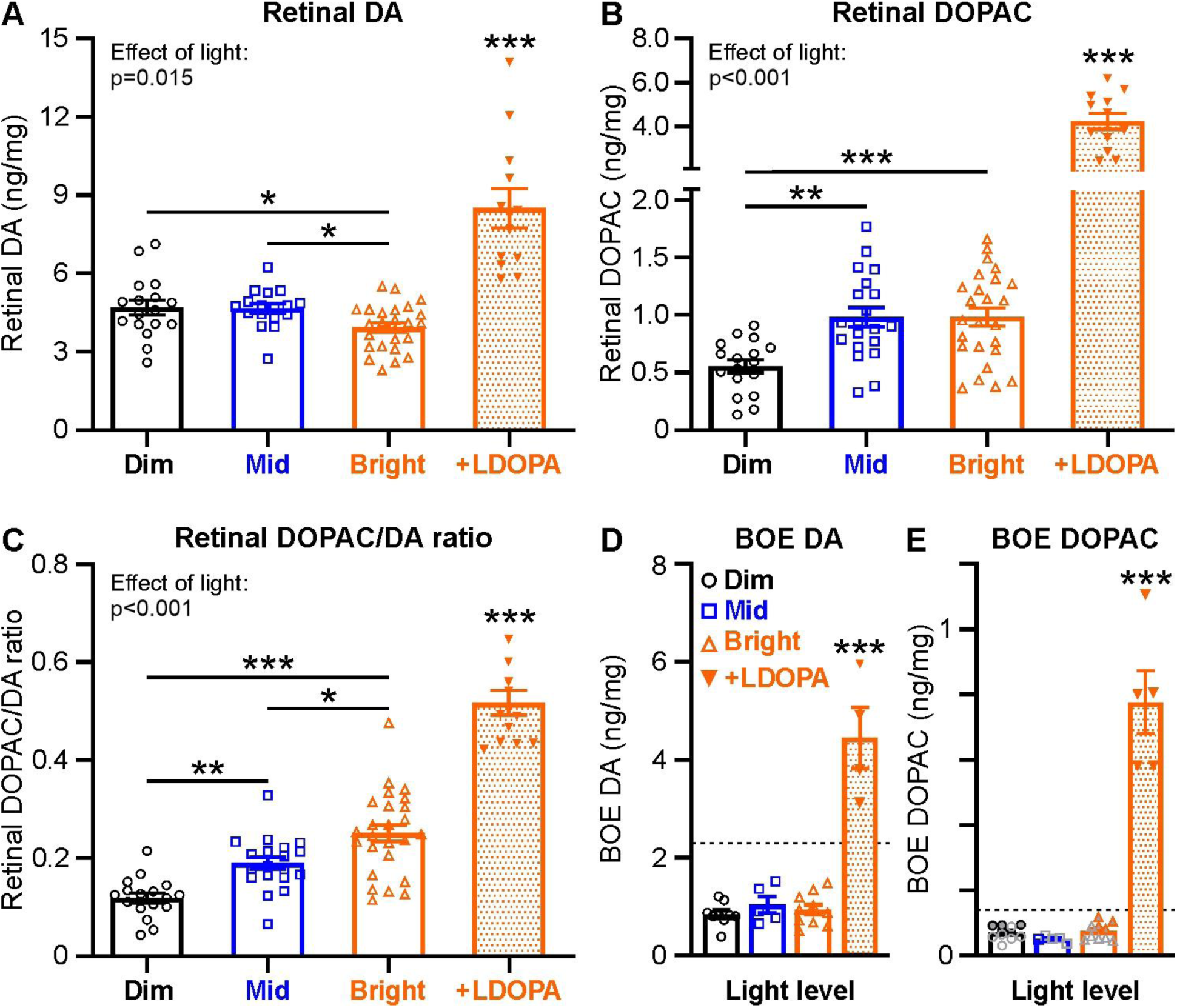
Brief exposure to bright ambient lighting increases DA turnover in the retina with no effect in the BOE. **A.** Average retinal DA levels, normalized to total retinal protein, after mice were exposed to Dim (black circles, n=17), Mid (blue squares, n=17), or Bright (orange triangles, n=25) ambient lighting for 5 minutes. As a positive control, a subset of mice exposed to Bright ambient lighting were also given IP injections of LDOPA (10 mg/kg) 10 minutes before light exposure (orange inverse triangles, n=12). **B-C.** As in panel A, but for average normalized retinal DOPAC levels (B, mid n=20) and the DOPAC/DA ratio, a measure of DA turnover (C, mid n=19). **D-E**. As in panels A-B, but for average BOE DA (D) and DOPAC (E) levels, normalized to total BOE protein (dim n=9, mid n=5, bright n=10, +LDOPA n=4 for DA and n=5 for DOPAC). Gray open symbols indicate that DOPAC measurements from both eyes were below the detection threshold and therefore arbitrarily set to 0.1 ng/ml before normalization and averaging. Symbols filled with gray indicate that the DOPAC measurement from only one eye was below the detection threshold and thus set to 0.1 ng/ml before normalizing and averaging values between the two eyes. Dashed lines denote the lowest measured value across all light levels for retinal DA and DOPAC, as comparison. Colors and symbols are the same in all subsequent figures unless otherwise specified. Statistical analysis was performed with one-way ANOVA with Tukey’s post hoc multiple comparisons test. Asterisks above LDOPA signifies significance compared to Dim, Mid, and Bright groups collectively. Data is shown as mean ± SEM.

Unlike in the retina, DA and DOPAC levels measured in the BOE were significantly lower and unaffected by ambient lighting (**Figure 2D-E**, one-way ANOVAs, effect of light, DA: p=0.527, DOPAC: p=0.158). The average BOE DA level across all light levels (0.90±0.06 ng/mg) was ∼61% lower than the lowest measured retinal DA level (Bright: 2.29 ng/mg) and was significantly lower than in LDOPA treated mice (4.44±0.63 ng/mg, p<0.001). Likewise, the average DOPAC level across all light levels in the BOE (0.07±0.005 ng/mg) was half that of the lowest DOPAC level in the retina (Dim: 0.14 ng/mg) and significantly lower than with LDOPA treatment (0.78±0.10 ng/mg p<0.001). Moreover, across light levels, 23 out of 24 BOE DOPAC measurements were below the HPLC detection threshold from at least one eye sample. From this data, we conclude that retinal DA turnover increases with light intensity, as shown in previous studies^17, 30^, but that this occurs rapidly after only 5 minutes of exposure to ambient lighting after dark adaptation. Importantly, we also find that DA signal dynamics do not change in the BOE where DA levels are low and DOPAC appears to not be present in any significant quantity, suggesting that DA signaling is restricted to the retina.

### atRA levels in the eye decrease with longer exposure to bright ambient lighting

Retinal atRA can continue to increase in the dark after a brief 10 minute light pulse^34^ but it is unclear how atRA levels change with increasing duration of exposure over a wide luminance range. Here, we exposed mice to Dim, Mid, and Bright ambient lighting for increasing amounts of time. Interestingly, we found that retinal atRA decreased as both ambient light intensity and exposure duration increased (**Figure 3A**, two-way ANOVA, duration x light interaction effect: p=0.036, effect of light: p=0.001, effect of duration: p=0.003). After any exposure duration, Bright light-exposed mice had significantly lower retinal atRA levels than in Dim light-exposed mice (5 min.: 46.23±1.27 vs 54.32±1.03 pmol/g, p<0.01; 15 min.: 41.74±1.42 vs 51.12±3.08 pmol/g, p<0.05; 60 min.: 32.89±1.85 vs 43.50±1.45 pmol/g, p<0.01). As exposure duration increased from 5 to 60 minutes, retinal atRA levels decreased in Dim and Bright ambient lighting (Dim: p<0.05, Bright: p<0.01), but there was no change in Mid ambient lighting (5 vs 60 min., 50.52±4.64 vs 52.09±1.28 pmol/g, p=0.826). Interestingly, atRA levels after 60 minutes of ambient light exposure were significantly higher under Mid lighting vs. either Dim or Bright lighting (Mid vs Dim / Bright: 52.09±1.28 vs 43.50±1.45 / 32.89±1.85 pmol/g; vs Dim p<0.01, vs Bright p<0.001). In the BOE, atRA levels similarly decreased with ambient light intensity (**Figure 3B**, two-way ANOVA effect of light: p=0.003) and exposure duration (two-way ANOVA effect of duration: p<0.001). There was also an effect of ambient light level and exposure duration on retinol (**Table 1**) and retinyl ester (**Table 2**) levels in the retina and BOE. However, the standard deviations of the data were very large, possibly due to batch effects in the dietary content of vitamin A that can significantly affect quantitative measurements of these retinoids and limit interpretation. Importantly, the batch effects present for retinol and retinyl esters did not affect atRA results. This observation is consistent with previous studies showing that it takes several generations of breeding on a diet to effect changes in atRA^50^ (e.g. within the first generation of a dietary change in vitamin A content, atRA is insensitive to changes in retinol). Overall, longer exposure to increased ambient light intensity reduces atRA levels in the retina and BOE. Surprisingly, however, in Mid light levels, longer exposure does not appear to influence retinal or BOE atRA.

**Figure 3.**
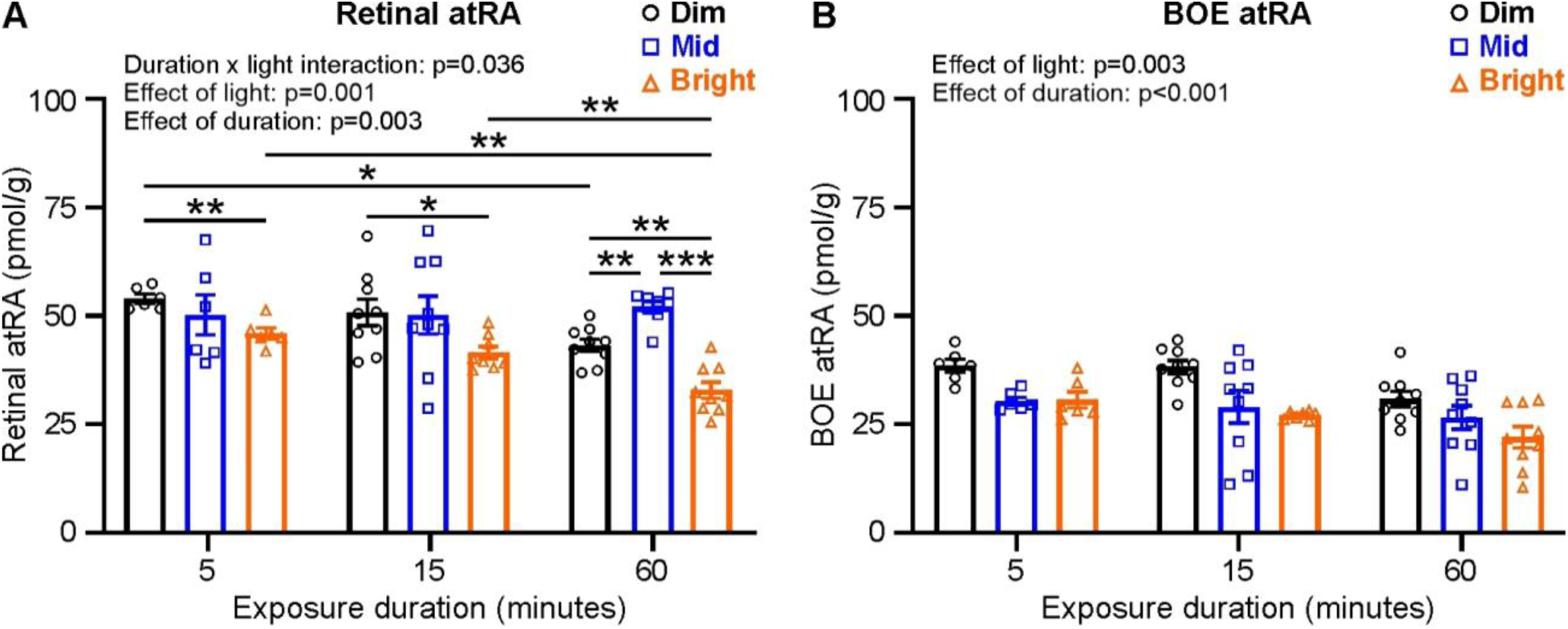
Ocular atRA levels decrease with ambient light intensity and exposure duration. **A**. Average retinal atRA levels, normalized to tissue weight, from mice exposed to Dim (black circles), Mid (blue squares), or Bright (orange triangles) ambient lighting for 5, 15, or 60 minutes (sample sizes in order of 5, 15, 60 minutes: Dim n=6, 9, 9; Mid n=6, 9, 8; Bright n=6, 8, 9). **B.** As in panel A, but for average normalized BOE atRA levels (15 min., Bright n=7; 60 min., Mid n=9,). Statistical analysis was performed with two-way repeated measures ANOVA with Tukey’s post hoc multiple comparisons test. Data is mean ± SEM.

**Table 1.**
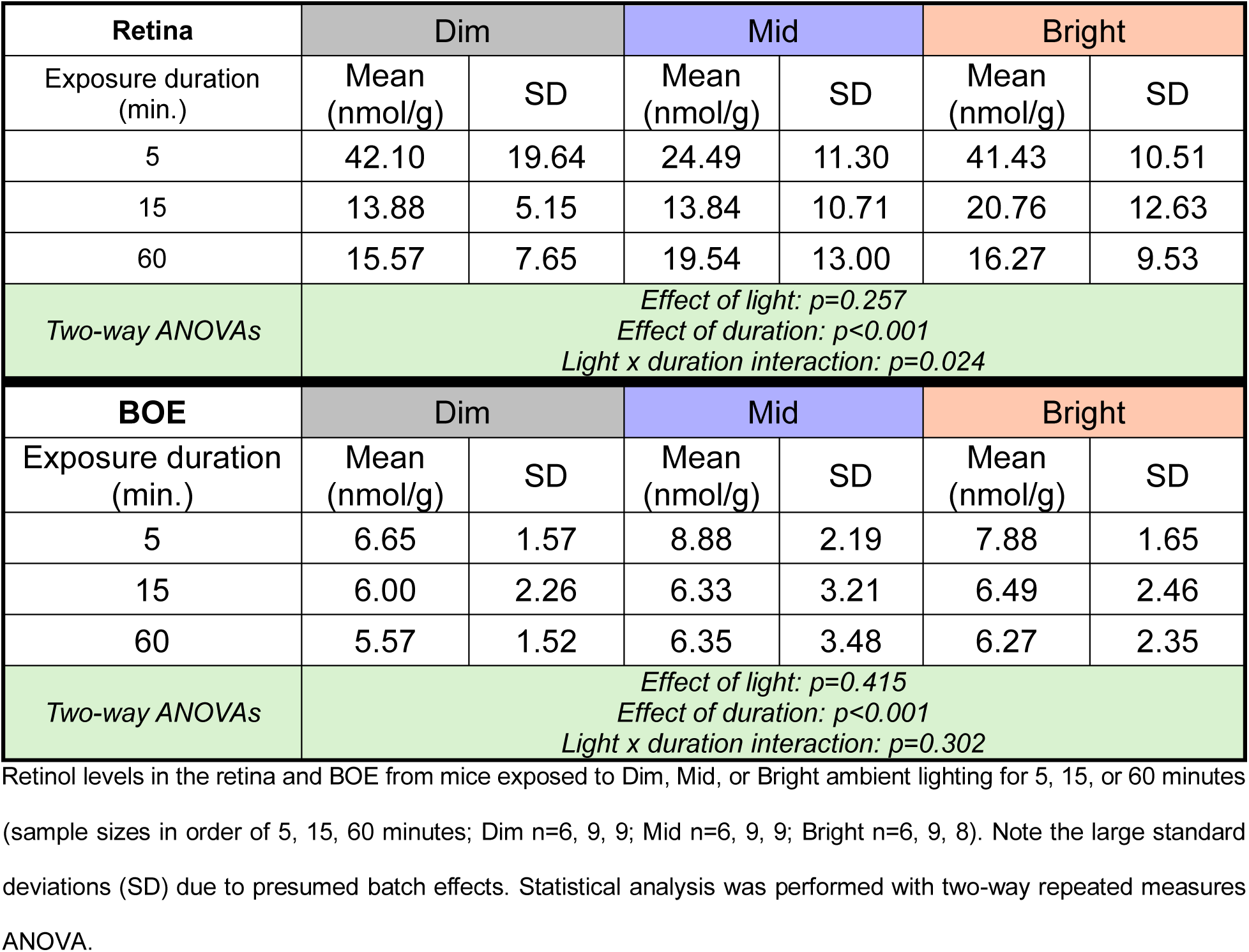
Retinol levels after ambient light exposure.

| **Retina** | **Dim** |  | **Mid** |  | **Bright** |  |
| --- | --- | --- | --- | --- | --- | --- |
| Exposure duration (min.) | Mean (nmol/g) | SD | Mean (nmol/g) | SD | Mean (nmol/g) | SD |
| 5 | 42.10 | 19.64 | 24.49 | 11.30 | 41.43 | 10.51 |
| 15 | 13.88 | 5.15 | 13.84 | 10.71 | 20.76 | 12.63 |
| 60 | 15.57 | 7.65 | 19.54 | 13.00 | 16.27 | 9.53 |
| Two-way ANOVAs | *Effect of light: $p=0.257$*<br>*Effect of duration: $p<0.001$*<br>*Light x duration interaction: $p=0.024$* |  |  |  |  |  |
| **BOE** | **Dim** |  | **Mid** |  | **Bright** |  |
| Exposure duration (min.) | Mean (nmol/g) | SD | Mean (nmol/g) | SD | Mean (nmol/g) | SD |
| 5 | 6.65 | 1.57 | 8.88 | 2.19 | 7.88 | 1.65 |
| 15 | 6.00 | 2.26 | 6.33 | 3.21 | 6.49 | 2.46 |
| 60 | 5.57 | 1.52 | 6.35 | 3.48 | 6.27 | 2.35 |
| Two-way ANOVAs | *Effect of light: $p=0.415$*<br>*Effect of duration: $p<0.001$*<br>*Light x duration interaction: $p=0.302$* |  |  |  |  |  |

Retinol levels in the retina and BOE from mice exposed to Dim, Mid, or Bright ambient lighting for 5, 15, or 60 minutes (sample sizes in order of 5, 15, 60 minutes; Dim n=6, 9, 9; Mid n=6, 9, 9; Bright n=6, 9, 8). Note the large standard deviations (SD) due to presumed batch effects. Statistical analysis was performed with two-way repeated measures ANOVA.

**Table 2.**
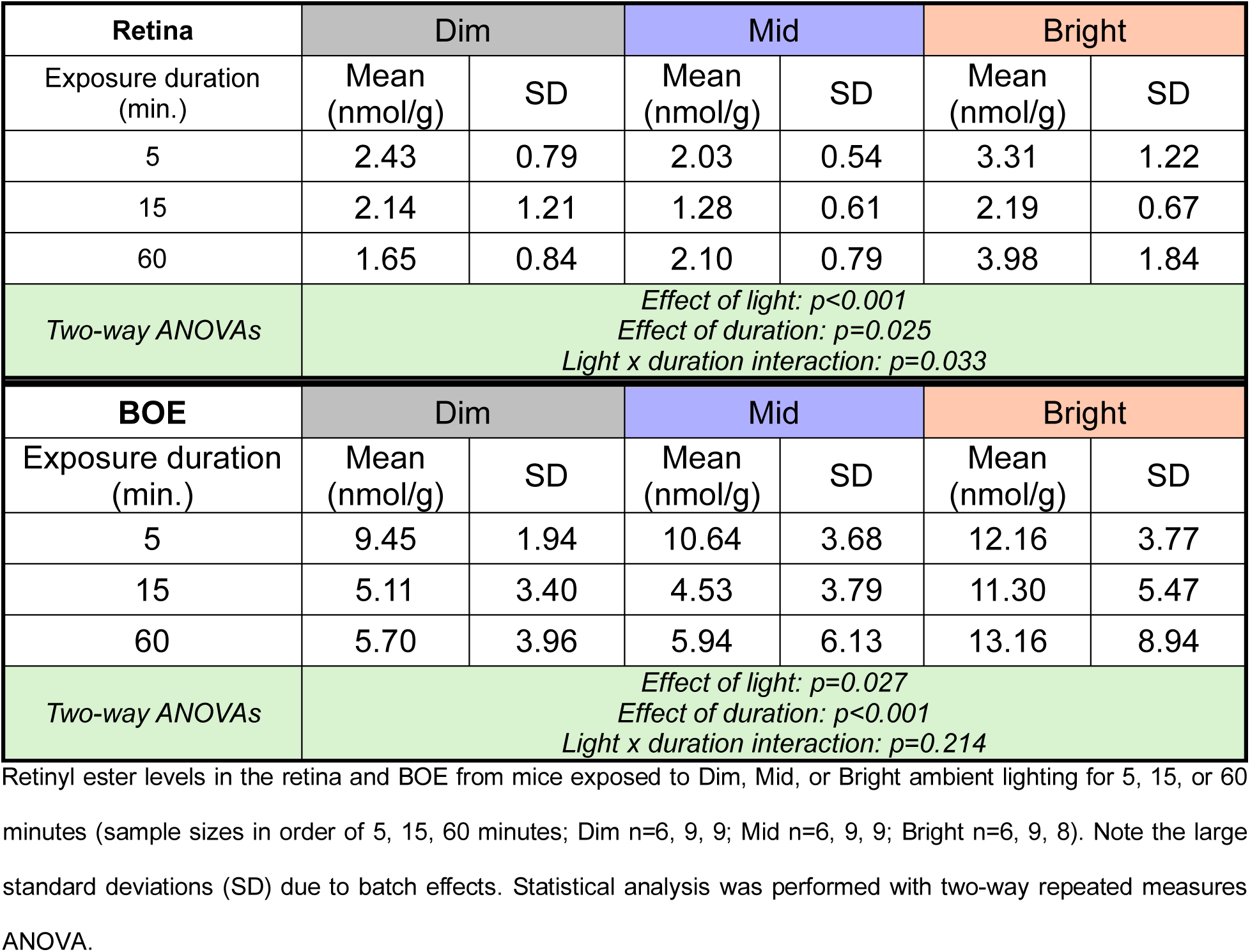
Retinyl ester levels after ambient light exposure.

### LDOPA transiently reduces retinal atRA levels which recover as light exposure increases

Retinal DA turnover and ocular atRA levels are oppositely affected by light intensity – DA turnover increases while atRA levels decrease when moving from Dim to Bright ambient conditions, and the effect on atRA is most pronounced with longer exposure time (**Figures 2 and** **3**). An intriguing question, therefore, is whether the effect of light intensity is mediated by direct interactions between DA signaling and atRA. To test this, IP injections of LDOPA (10 mg/kg) were given to mice 10 minutes before exposure to Mid ambient lighting. Mid ambient lighting was chosen since there was no change in atRA levels at this intensity (**Figure 3**). Surprisingly, LDOPA treatment had a significant effect on retinal atRA (**Figure 4A**, two-way ANOVA, duration x treatment interaction effect: p=0.038, effect of treatment: p<0.001, effect of duration: p=0.043). After 5 minutes of Mid light exposure, LDOPA-treated mice had a pronounced reduction of almost 50% in retinal atRA compared to saline-treated mice (23.64±0.73 vs 40.40±2.24 pmol/g, p<0.001). This reduction was attenuated but still present after 15 minutes (29.64±3.79 vs 38.71±0.74 pmol/g, p<0.05) and recovered to control levels after 60 minutes (35.85±1.24 vs 40.23±3.17 pmol/g, p=0.230; +LDOPA 5 vs 60 min.: p<0.01). In contrast, there was no effect of LDOPA on atRA levels in the BOE (**Figure 4B**, two-way ANOVA, effect of treatment: p=0.486, effect of duration: p=0.766). Unlike retinoid measurements in different ambient lighting, there was no effect of LDOPA on retinol (**Table 3**) or retinyl esters (**Table 4**) in the retina or BOE. While batch effects and large standard deviations limit interpretation as with the previous data, they did not impact the relationship between LDOPA and atRA. Overall, these data suggest a direct influence of DA on retinal atRA.

**Figure 4.**
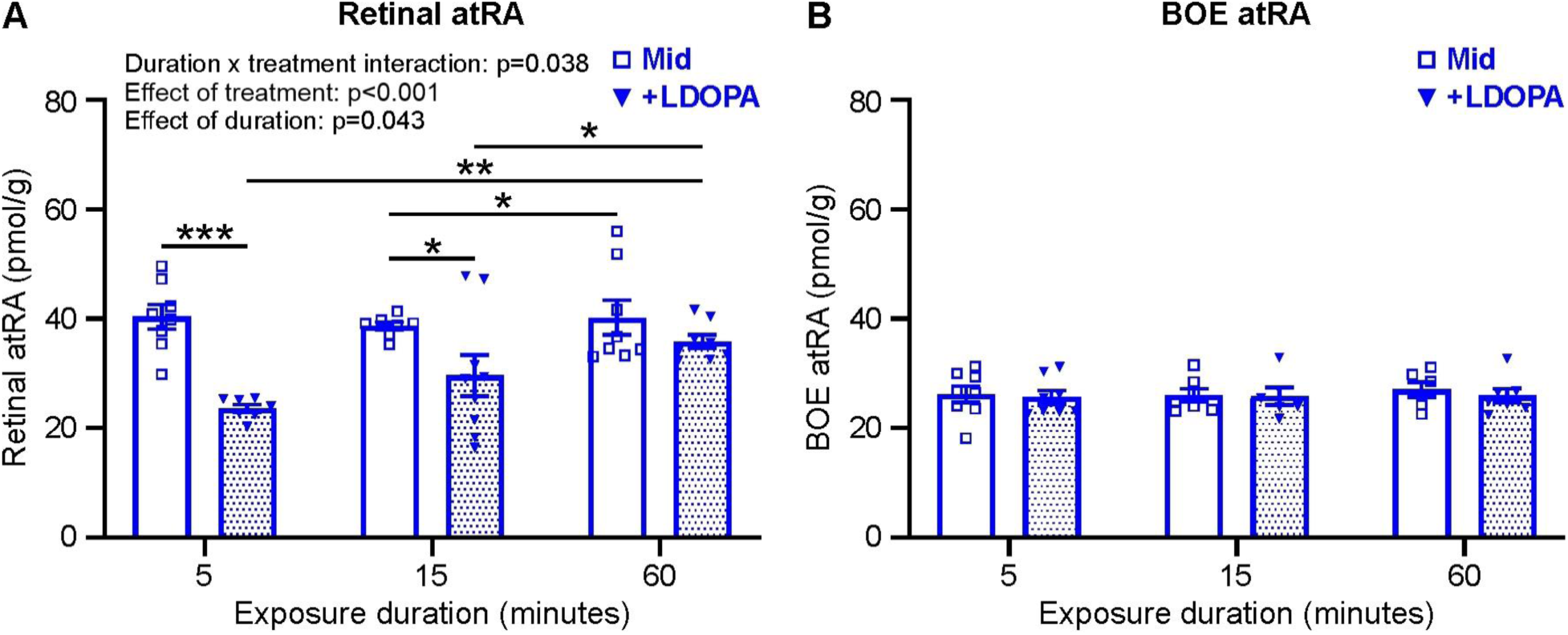
Retinal atRA levels are reduced with LDOPA treatment. **A.** Average retinal atRA levels, normalized to tissue weight, from mice exposed to Mid ambient lighting that received IP injections of saline alone (blue squares) or LDOPA dissolved in saline (blue inverse triangle, 10 mg/kg) 10 minutes before light exposure for 5, 15, or 60 minutes (sample sizes in order of 5, 15, 60 minutes: Mid n=8, 7, 8; +LDOPA n=7, 9, 8). **B.** Same as in A but for average normalized BOE atRA levels (sample sizes in order of 5, 15, 60 minutes: Mid n=8, 7, 6; +LDOPA n=8, 6, 7). Statistical analysis was performed with two-way repeated measures ANOVA with Tukey’s post hoc multiple comparisons test. Data is mean ± SEM.

**Table 3.** Retinol levels after LDOPA treatment.

| **Retina** | **Mid + saline** |  | **Mid + LDOPA** |  |
| --- | --- | --- | --- | --- |
| Exposure duration (min.) | Mean (nmol/g) | SD | Mean (nmol/g) | SD |
| 5 | 25.03 | 15.77 | 25.66 | 16.56 |
| 15 | 20.96 | 15.36 | 21.52 | 15.16 |
| 60 | 20.08 | 22.70 | 13.52 | 9.17 |
| *Two-way ANOVAs* | *Effect of treatment: p=0.758*<br>*Effect of duration: p=0.029*<br>*Treatment x duration interaction: p=0.336* |  |  |  |
| **BOE** | **Mid + saline** |  | **Mid + LDOPA** |  |
| Exposure duration (min.) | Mean (nmol/g) | SD | Mean (nmol/g) | SD |
| 5 | 5.29 | 1.46 | 5.42 | 0.75 |
| 15 | 5.29 | 0.97 | 5.32 | 1.35 |
| 60 | 5.89 | 2.00 | 5.57 | 1.13 |
| *Two-way ANOVAs* | *Effect of treatment: p=0.838*<br>*Effect of duration: p=0.530*<br>*Treatment x duration interaction: p=0.828* |  |  |  |

Retinol levels in the retina and BOE from mice given IP injections of saline along or saline + LDOPA (10 mg/kg) 10 minutes before exposure to Mid ambient lighting for 5, 15, or 60 minutes (sample sizes in order of 5, 15, 60 minutes; retina: saline n=8, 9, 9; LDOPA n=8, 9, 8; BOE: saline n=8, 7, 7; LDOPA n=8, 7, 6). Note the large standard deviations (SD) due to batch effects. Statistical analysis was performed with two-way repeated measures ANOVA.

**Table 4.** Retinyl ester levels after LDOPA treatment.

| **Retina** | **Mid + saline** |  | **Mid + LDOPA** |  |
| --- | --- | --- | --- | --- |
| Exposure duration (min.) | Mean (nmol/g) | SD | Mean (nmol/g) | SD |
| 5 | 2.40 | 1.32 | 2.11 | 1.33 |
| 15 | 1.09 | 0.84 | 1.21 | 1.10 |
| 60 | 1.32 | 1.04 | 1.19 | 1.04 |
| *Two-way ANOVAs* | *Effect of treatment: p=0.871*<br>*Effect of duration: p=0.013*<br>*Treatment x duration interaction: p=0.866* |  |  |  |
| **BOE** | **Mid + saline** |  | **Mid + LDOPA** |  |
| Exposure duration (min.) | Mean (nmol/g) | SD | Mean (nmol/g) | SD |
| 5 | 5.92 | 3.47 | 5.86 | 2.03 |
| 15 | 6.70 | 3.50 | 6.07 | 2.05 |
| 60 | 6.26 | 2.44 | 5.36 | 2.32 |
| *Two-way ANOVAs* | *Effect of treatment: p=0.524*<br>*Effect of duration: p=0.806*<br>*Treatment x duration interaction: p=0.861* |  |  |  |

Retinyl ester levels in the retina and BOE from mice given IP injections of saline along or saline + LDOPA (10 mg/kg) 10 minutes before exposure to Mid ambient lighting for 5, 15, or 60 minutes (sample sizes in order of 5, 15, 60 minutes; retina: saline n=8, 9, 9; LDOPA n=8, 9, 8; BOE: saline n=8, 7, 7; LDOPA n=8, 7, 6). Note the large standard deviations (SD) due to batch effects. Statistical analysis was performed with two-way repeated measures ANOVA.

### Exogenous atRA treatment does not affect ocular DA turnover

Systemic LDOPA treatment affects retinal, but not BOE, levels of atRA (**Figure 4**). We next asked whether the opposite was true, i.e. do atRA levels affect DA in the eye? To address this question, mice were trained to voluntarily ingest 1% atRA (25 mg/kg) or vehicle sugar pellets two hours before ambient light exposure. Two hours after atRA ingestion, retinal atRA levels were significantly elevated (more than 8-fold) compared to retinas from sugar-fed mice (385.6±123.4 vs 46.5±8.8 pmol/g, n=2, data not shown). Interestingly, retinal DA levels were unaffected by atRA treatment regardless of ambient light intensity (**Figure 5A**, two-way ANOVA, effect of treatment: p=0.959). However, note that retinal DA significantly decreased with brighter ambient exposure as in **Figure 2** (effect of light: p=0.003). Likewise, there was no significant effect of atRA treatment on retinal DOPAC levels across ambient light conditions (**Figure 5B**, two-way ANOVA, effect of treatment: p=0.341) though the DOPAC level in Mid light was on average lower with atRA treatment than control (atRA vs ctrl, 0.41±0.03 vs 0.58±0.08 ng/mg, p=0.138, Mann-Whitney test). As a result, there was little difference between Dim and Mid light levels in atRA fed mice, unlike in control mice, despite an overall significant increase in DOPAC with brighter ambient light exposure (effect of light: p<0.001). Expectedly, the lack of significant differences on DA and DOPAC led to no difference in DA turnover (DOPAC/DA ratio) between atRA-fed and control mice (**Figure 5C**, two-way ANOVA, effect of treatment: p=0.641), although DA turnover increased with light intensity as seen before (effect of light: p<0.001, **Figure 2**). Lastly, there was no effect of atRA treatment on DA or DOPAC in the BOE (**Figure 5D-E**), and the average across ambient light levels and treatments was lower than the lowest retinal DA or DOPAC measurements (dashed lines in panels D-E). Therefore, it appears that increased atRA does not affect ocular DA turnover.

**Figure 5.**
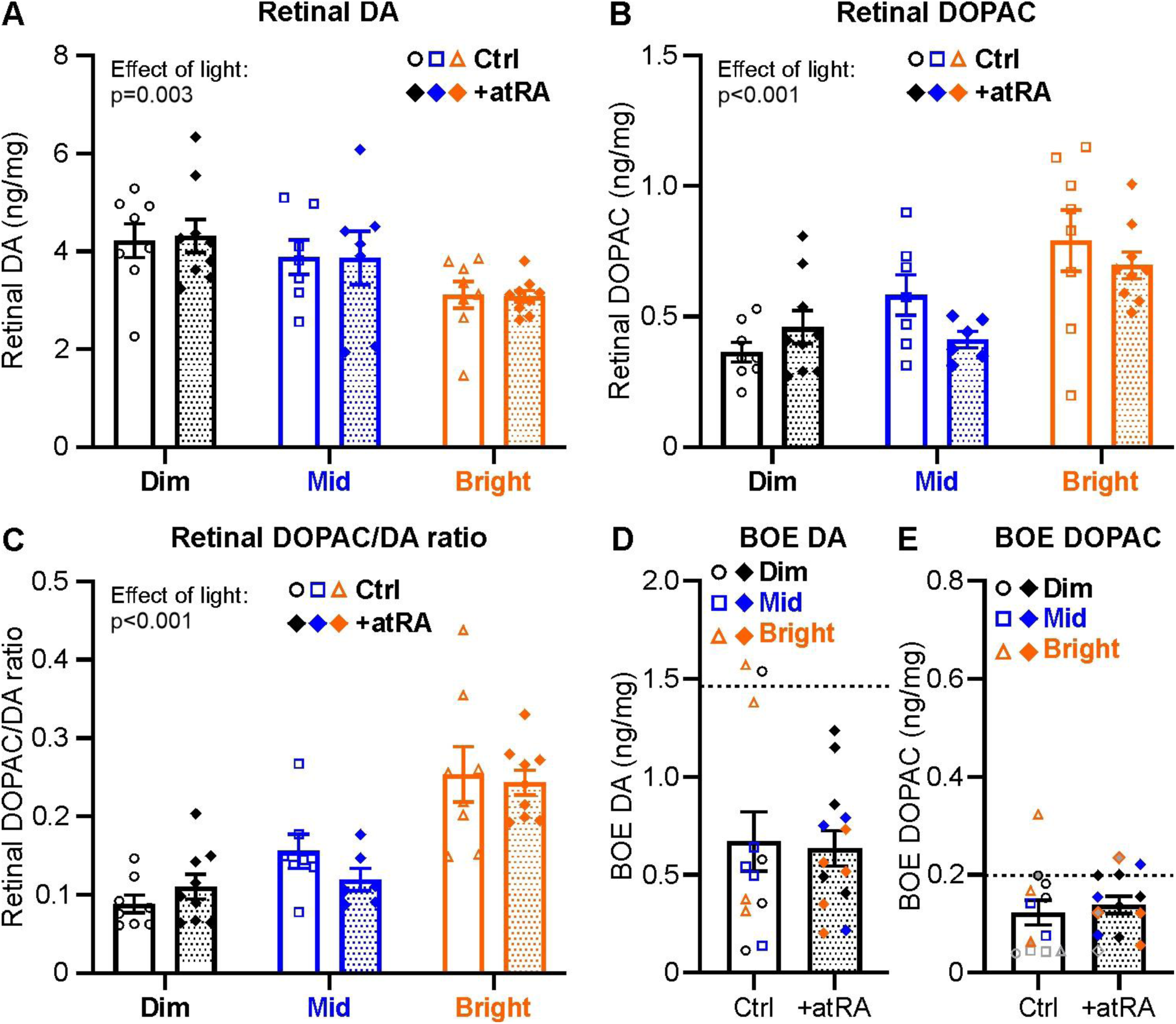
Ocular DA turnover is not influenced by exogenous atRA treatment. **A.** Average retinal DA levels, normalized to total retinal protein, in mice exposed to Dim (black circles, n=8), Mid (blue squares, n=7), or Bright (orange triangles, n=8) ambient lighting for five minutes after voluntary ingestion of 1% atRA sugar pellets or vehicle pellets (solid diamonds, 25 mg/kg, Dim n=9, Mid n=7, Bright n=9) two hours before light exposure. **B-C.** As in panel A, but for average normalized retinal DOPAC levels (B, Mid + atRA n=6) or the DOPAC/DA ratio (C, Mid + atRA n=6). **D-E.** As in panels A-B, but for average BOE DA (D, ctrl n=12, +atRA n=13) and DOPAC (E, ctrl n=12, +atRA n=13) across all ambient light levels, normalized to total BOE protein. Gray open symbols indicate that DOPAC measurements from both eyes were below the detection threshold and therefore arbitrarily set to 0.1 ng/ml before normalization and averaging. Symbols filled with gray indicate that the DOPAC measurement from only one eye was below the detection threshold and thus set to 0.1 ng/ml before normalizing and averaging values between the two eyes. Dashed lines denote the lowest measured value across all light levels for retinal DA and DOPAC as comparison. Statistical analysis was performed with two-way repeated measures ANOVA with Tukey’s post hoc multiple comparisons test. Data is mean ± SEM.

## Discussion

Our study supports three main conclusions: 1) ocular atRA levels decrease as ambient illumination or exposure duration increases; 2) modulation of ocular atRA and retinal DA signaling by ambient illumination occurs rapidly; and 3) DA has an acute inhibitory effect on retinal atRA. An increase in total retinal illumination, either through increased brightness or prolonged exposure, reduced retinal and BOE atRA (**Figure 3**). Interestingly, this effect was restricted to Dim and Bright ambient lighting, as there was no effect of exposure duration in Mid ambient lighting. In contrast, brief exposure to increased light intensity was sufficient to increase DA turnover (**Figure 2**) similar to more prolonged exposure durations demonstrated in previous work^17, 30^. Furthermore, increasing DA through systemic administration of LDOPA transiently reduced retinal atRA whereas systemic administration of atRA had no effect on ocular DA (**Figures 4 and 5**). Overall, these data suggest interactions between ambient light, atRA, and DA that may contribute to a retino-scleral signaling cascade (**Figure 6**). However, it is unclear if the effect of bright light on atRA occurs directly or through increased DA signaling and whether reduced atRA in the BOE with bright light is mediated by local signaling or is a consequence of signaling from the retina. Lastly, DA signaling appears to be limited in the BOE.

**Figure 6.**
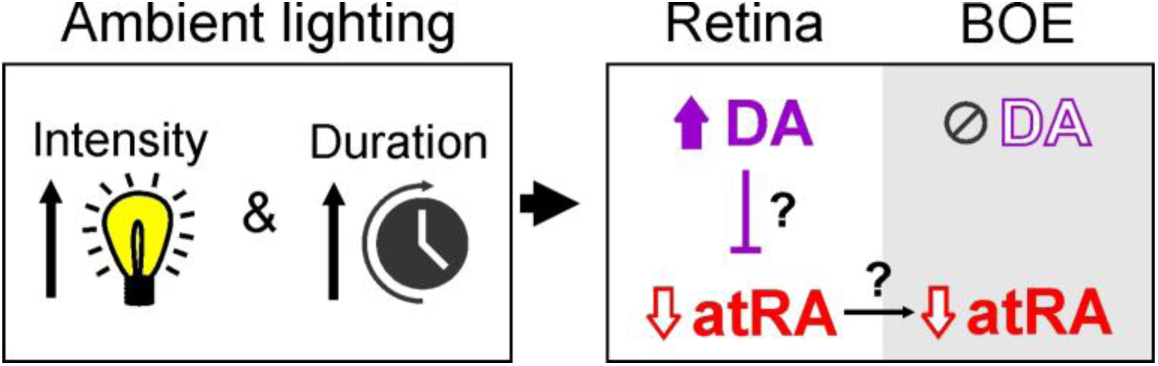
Proposed signaling cascade involving interactions between light exposure, DA signaling (purple), and atRA (red) in the retina and the BOE. As ambient light intensity and duration increase up to 60 minutes, retinal DA turnover increases while retinal and BOE atRA decreases. Increased DA also reduces retinal atRA. However, it is unclear if the reduction in retinal atRA with increased ambient lighting is driven by higher retinal DA turnover. Additionally, it is not known whether reduced BOE atRA is due to local factors or is influenced by reductions in retinal atRA.

### Ambient light conditions have intensity- and temporal-dependent effects on ocular atRA

atRA, a derivative of vitamin A and a critical physiological signaling molecule in eye development^51^, is ultimately synthesized from retinol primarily by the retinal pigment epithelium (RPE)^34^ but also by other retinal^35–38^ and choroidal cells^33, 39^. While retinal and choroidal atRA can be modulated by visual blur^25, 33, 52–54^, only one study has investigated atRA’s dependence on ambient light level^34^. McCaffrey and colleagues found that retinal atRA release in dark-adapted mice increased after light exposure compared to darkness^34^. They analyzed supernatant from cultured retinas after a 10-minute light pulse followed by dark-incubation and from eye cup (BOE + retina) supernatant sampled over two hours of light exposure. Interestingly, after an initial increase in atRA with light exposure, they found that atRA levels tended to decrease with exposure duration. Our results support these findings and provide more specific and definitive information about the effects of light on ocular atRA. First, we find a significant temporal interaction between light and retinal atRA: as exposure duration increases, atRA levels decrease (**Figure 3**, note that we did not compare atRA levels to a dark-adapted state). Moreover, McCaffrey and colleagues did not test multiple ambient intensities, using only a light plate of unspecified irradiance under “bright room light”. Though light exposure increases retinal atRA, we demonstrate that the increase depends on ambient intensity and on which ocular tissue is being assayed (**Figure 3**). Initially, after 5 minutes of light, retinal atRA levels followed a monotonic intensity-dose relationship (Dim>Mid>Bright). However, as exposure duration increased, retinal and BOE atRA levels in both Dim and Bright ambient lighting decreased, while atRA levels from retinas exposed to Mid ambient lighting did not change. This differential modulation resulted in retinal atRA being highest after 60 minutes of Mid light exposure. Intriguingly, the Mid light level (59 lux) is just past the mesopic-photopic visual threshold^55^. Since rod photoreceptors can be active at brighter intensities than previously thought^56^, both rod and cone pathways may be active at this light level but sub-optimally active (i.e. ambient lighting is too bright for rods and not bright enough for cones for optimal stimulation), leading to relatively higher atRA levels. In contrast, the Dim ambient intensity (1 lux) is just below the mesopic regime^55^ and as exposure duration increases, rod pathway activation may increase, reducing atRA. Similarly, in Bright ambient light (12,000 lux) cones are strongly activated, reducing atRA. Unlike in the retina, atRA levels in the BOE followed a monotonic relationship (Dim>Mid>Bright) regardless of exposure duration. Ultimately, mechanistic studies are required to test photoreceptor pathway contribution to ocular atRA levels and to determine whether retinal atRA influences the atRA in the BOE.

### Modulation of ocular DA by ambient light occurs within minutes

Ocular DA is released from retinal DACs^29^ and activates receptors located throughout the retina and the RPE^28, 29^. After release, DA is transported back into cells and metabolized to DOPAC^57, 58^, with the ratio of DOPAC to DA reflecting DA turnover^59^. DA release, synthesis, and metabolism increase with brightness^60, 61^. In a series of recent studies, DA signaling was further investigated over a range of ambient light levels. Perez-Fernandez and colleagues demonstrated that DA turnover was low and stable throughout the scotopic (rod only) and mesopic ranges (rod + cone) but increased within the photopic range (cone only, ∼400 lux)^30^. DA turnover was measured from whole retinas or isolated eye cup supernatant after mice or eye cups received a light pulse (15 minutes – 1 hour). They also found that DA rapidly peaked within minutes of light exposure before reaching a steady-state by 15 minutes. Consistent with these findings, we demonstrate that after only 5 minutes of light exposure, whole retina levels of DA and DOPAC are significantly affected as a function of ambient brightness, leading to greater DA turnover with brighter ambient light (**Figure 2**). This was not true in the BOE, where DA levels were low and unaffected by ambient lighting, and DOPAC was minimal (**Figure 2**), suggesting that DA does not play a major role exterior to the retina and RPE.

Moreover, DA dynamics appear to depend on the time-course of ambient light exposure. The effect of ambient light on DA tunrover that we find after only 5 minutes (**Figure 2**) is maintained after either a more prolonged 3-hour light pulse or after rearing mice in different light levels for 2 weeks^17^. Interestingly, while DOPAC levels consistently increase after exposure to brighter ambient lighting regardless of exposure duration, it appears that DA levels only decrease with more acute exposure, since rearing mice in different ambient lighting had no effect^17^. This could reflect adaptation of DA release, which may reach a maximal output, while DA turnover and metabolism continue to increase^28^. Furthermore, although the Dim and Mid light levels used in this study were brighter than the levels used previously^17^, the overall effect of ambient light remains consistent across exposure durations between studies.

Lastly, in this study we analyzed whole retinas, an approach which captures both released and un-released DA and DOPAC. However, vitreal DOPAC is proposed to be a surrogate for retinal DA release^59^, and DOPAC levels in collected vitreous samples increased after exposure to bright light in chickens^62–64^. Of note, DOPAC concentration in the vitreous continues to rise over the light phase^64^, suggesting that vitreal DOPAC measurements may reflect more accumulation rather than temporally accurate release. Overall, we demonstrate that the effect of different ambient lighting on retinal DA occurs rapidly, suggesting that even brief changes in ambient illumination, such as turning on or off indoor lights, or looking around a visual scene, can alter DA signaling dynamics. One remaining question however, is whether the temporal profile of DA turnover to reach steady-state levels after 15 minutes changes with ambient light level, as only the brightest, photopic level was tested by Perez-Fernandez et al^30^.

### Transient inhibition of retinal atRA by DA

Excitingly, we found that systemic administration of LDOPA, the DA precursor, resulted in reduced retinal atRA after 5 minutes of mid light exposure which recovered to control levels by 60 minutes (**Figure 4**). However, systemic administration of atRA did not affect retinal DA turnover (**Figure 5**). These data suggest a unidirectional acute inhibition of atRA by DA. Two important questions arise from this novel finding: 1) Is the increase in DA turnover with bright ambient lighting sufficient to elicit a transient reduction in atRA as seen with LDOPA? and 2) is the reduction of atRA in brighter ambient lighting independent of increased DA turnover?

To address the first of these questions, we note that LDOPA administration almost doubled DA turnover after mice were exposed to Bright ambient light (**Figure 2**). Therefore, we assume that LDOPA also significantly increased DA turnover in the Mid ambient lighting that we used. While we do see an inverse correlation between DA and atRA with ambient light level, it is still unclear if increased DA turnover within physiological levels, such as the doubling of DA turnover from Dim to Bright ambient lighting (**Figure 2**), would elicit the same atRA response as LDOPA. Regarding the second question, we have shown that retinal atRA is reduced following two independent manipulations: ambient light and LDOPA. Since DA turnover increases in Bright light, DA signaling could be the mechanism underlying the effect of ambient lighting on atRA rather than direct effects of light itself (**Figure 6**). Nonetheless, our findings raise important implications of the DA-atRA interaction for visual processing, eye growth, and myopia.

### Implications of ambient lighting on atRA and DA in myopia development

Outdoor environments and brighter light protect against myopia onset in children and attenuate experimental myopia in animal models^3, 8–22^, although Dim light was also recently associated with less myopia in children and mice while Mid light levels were correlated with the most myopia^3, 17^. Additionally, both atRA and DA are associated with opposing endpoints in eye growth and development: DA as a ‘stop’ signal in myopic eye growth^23, 24^ and atRA as a ‘go’ signal^25–27^. Activation of DA signaling attenuates induced myopia in animal models^17, 65–70^, depletion of DA signaling can lead to spontaneous myopia in mice^71^, and DA or DOAPC is reduced in myopic eyes^23, 72–77^ (though see^78–81^ showing no effect in mice). In contrast, administration of atRA induces myopic eye growth^26, 53, 82^, inhibition of atRA signaling protects against myopia development^54, 83^, and atRA levels are increased in myopic eyes^25, 46, 52, 53, 83, 84^. Interestingly, we find that both Dim and Bright ambient lighting inhibited retinal atRA more than Mid ambient lighting after 60 minutes of exposure (**Figure 3**). On the other hand, DA turnover was highest in Bright conditions and lowest in Dim conditions. Therefore, retinal atRA was higher and DA turnover lower in light conditions that are implicated in myopia (i.e. indoor light levels). Moreover, retinal atRA was lower and DA turnover higher in Bright lighting which is associated with reduced myopia. Interestingly, Dim lighting, which was associated with reduced myopia in mice^17^, yielded higher atRA and lower DA turnover (**Figures 2 and** **3**), though it is still unclear the mechanisms that underly the protective effect of dim light and whether this finding is common across experimental models. Nonetheless, our findings support a ‘push-pull’ effect of ambient lighting on atRA and DA and match well their associations with myopia.

Importantly, DA and atRA receptor expression are localized to different posterior eye wall compartments: DA to the retina and RPE and atRA throughout the eye (retina, RPE, choroid, and sclera)^27, 29, 51, 85^. Therefore, a major question is how ambient light signals are translated through each tissue to ultimately influence sclera remodeling which occurs during eye growth and myopic axial elongation^26, 33, 39, 46, 86^. Our present study provides more information about potential roles of DA and atRA in this retino-scleral signaling cascade. Despite the significant effects of DA in myopia, DA is not likely a scleral signal and from our results, DA (and especially DOPAC) appear to be limited in the BOE (**Figure 2**). An intriguing theory is that DA signaling influences retinal atRA which may then act directly or indirectly on the posterior ocular layers (e.g. RPE, choroid, and sclera). While it is unclear if and how atRA from the retina could reach these layers, choroidal atRA can diffuse to and ready accumulate in the sclera^39^. While the exact source of sclera atRA is unknown (i.e. retinal vs choroidal), we found that retinal levels of atRA were higher than in the BOE, which includes the choroid, and that retinal atRA had the most change with light exposure and LDOPA treatment (**Figures 3 and** **4**). In general, the effect of ambient lighting on atRA was similar between the retina and BOE, further suggesting that atRA in the BOE is produced in the retina. Furthermore, the RPE could potentially participate in regulation of atRA transport to the sclera since DA receptor activation on RPE cells modulates ionic and fluid flow to the choroid^87–90^. This change in fluid transport could potentially facilitate transport of atRA, since RPE fluid transport is modulated in myopic animals^91–93^; however, these links require more direct testing to evaluate this hypothesized transport mechanism.

In summary, we find pronounced effects of ambient light intensity and duration on ocular DA and atRA levels in an ocular tissue-dependent manner (retina vs BOE). Additionally, we find an intriguing interaction between DA and atRA – namely, increased DA signaling transiently reduces retinal atRA. Therefore, we propose a retino-scleral signaling cascade that involves visual input (i.e. ambient lighting) modulating retinal DA levels which mediate atRA levels in the BOE, (**Figure 6**). However, further mechanistic investigations are needed to reveal direct connections between ambient lighting, DA, and atRA.

## Acknowledgements

We would like to thank Dr. Rong Fu and the Emory HPLC Bioanalytical Core. This study was supported in part by the Emory HPLC Bioanalytical Core (EHBC), which is subsidized by the Emory University School of Medicine and is one of the Emory Integrated Core Facilities. Additional support was provided by the Georgia Clinical & Translational Science Alliance of the National Institutes of Health under Award Number UL1TR002378. The content is solely the responsibility of the authors and does not necessarily reflect the official views of the National Institutes of Health.

